# Empowering STEM Students: A University-wide Mentorship Program Fostering Retention and Belonging

**DOI:** 10.1101/2024.04.02.587784

**Authors:** Sumedha Ravishankar, Tara C.J. Spencer-Drakes, Ivy H. Fernandes, Megan I. Hayes, Shelby Coopwood, Ilya Spencer, Sonya E. Neal

**Affiliations:** School of Biological Sciences, Department of Cell and Developmental Biology, University of California San Diego, La Jolla, CA 92093, USA; Howard Hughes Medical Institute, Chevy Chase, MD 20815, USA; Antigua and Barbuda International Institute of Technology, Piggotts, Antigua & Barbuda

**Keywords:** Mentorship, BUMMP, STEM, Retention, Professional Development, Research

## Abstract

In the face of a challenging climate that is resistant to Diversity, Equity, and Inclusion (DEI) efforts, there is a critical need for a support structure for retaining students, particularly those from historically excluded groups (HEGs), in STEM. The Biology Undergraduate and Master’s Mentorship Program (BUMMP) embodies this commitment to fostering scientific identity, efficacy, and sense of belonging for 1^st^ generation and historically underserved undergraduate and master’s students at UC San Diego. The mission of BUMMP is to cultivate a sense of belonging, instill confidence, and nurture a strong scientific identity amongst all of its participants. At its core, the three pillars of BUMMP are: 1) mentorship, 2) professional development, and 3) research. Quality mentorship is provided where students receive personal guidance from faculty, graduate students, postdocs, and industry leaders in navigating their career pathways. Complementing mentorship, BUMMP provides paid research opportunities and prioritizes professional development by offering workshops designed to enhance students’ professional skills. These three pillars form the backbone of BUMMP, empowering students from all backgrounds and ensuring their retention and persistence in STEM. So far, we’ve served over 1,350 mentees, collaborated with 809 mentors, and had over 180 mentees actively engaged in BUMMP-sponsored research activities. Overall, BUMMP’s expansive efforts have made a tremendous impact at UC San Diego and will continue to foster a community of future leaders who will be prepared to make meaningful contributions to the scientific community and beyond.

## Introduction

Systemic oppression and inequality in academic institutions are preventing undergraduate students from historically excluded groups (HEG) from flourishing in higher education in STEM. National studies suggest that Latinx and Black students choose a STEM major at the same rate as White students(Riegle-Crumb et al. 2019; Lewis-McCoy 2020). However, historically excluded students were more likely to switch out of their STEM major or discontinue their education(Lewis-McCoy, 2020). For example, due to a deep-rooted history of discriminatory practices in the United States, historically excluded students are more likely to attend under-resourced K-12 schools, making them less academically prepared compared to their peers from well-resourced schools (Bryant 2015). Upon entering higher education, HEG students face discrimination in an unwelcoming campus climate and lack relatable role models amongst faculty (Gladstone and Cimpian 2021). This has led to lower retention rates for HEG students, who grapple with diminished belonging, weakened STEM identity, and reduced science efficacy. The pandemic has exacerbated these existing disparities, disproportionately affecting HEG students who entered the crisis with the greatest needs and fewest opportunities(Gopalan et al., 2022; Herres et al., 2023). Furthermore, the Supreme Court’s decision against affirmative action has threatened DEI efforts in academic institutions, resulting in scaled-back initiatives (Noller et al., 2024; Tedesco, 2005). Hence, current initiatives to increase and retain HEG student representation in STEM higher education are needed more than ever.

Immediate action is needed for nationwide and university-level reforms to counteract the declining climate and enhance retention rates among HEG students. A study by Estrada et al. revealed a positive correlation between quality mentorship and research experience with an undergraduate student’s sense of belonging, scientific identity, and science efficacy(Estrada et al., 2018). This underscores the pivotal role mentors and researchers can play in shaping the future of HEG students and boosting STEM retention. Implementing a structured mentorship program alongside offering research experiences could significantly enhance HEG retention rates, especially when “scaled up” to reach more students. The Biology Undergraduate and Mentor’s Program (BUMMP) at University of California San Diego (UC San Diego), provides a promising model for increasing retention of HEG undergraduate and master’s students and preparing them to succeed in graduate and professional programs. Despite being only 4 years old, BUMMP has served over 1,350 students to date, setting it apart as a pioneering initiative with unmatched scale in major universities. This paper describes the three major pillars of BUMMP: 1) mentorship, 2) professional development, and 3) research (Figure 1). In addition, this paper examines how each component of BUMMP has shaped students’ sense of belonging, science identity, and science efficacy, which has been shown to predict retention and persistence to pursue a STEM career. Finally, we explore the key factors contributing to BUMMP’s success, emphasizing core values and the program’s foundation rooted in the lived experiences of its organizers.

**Figure 1:**
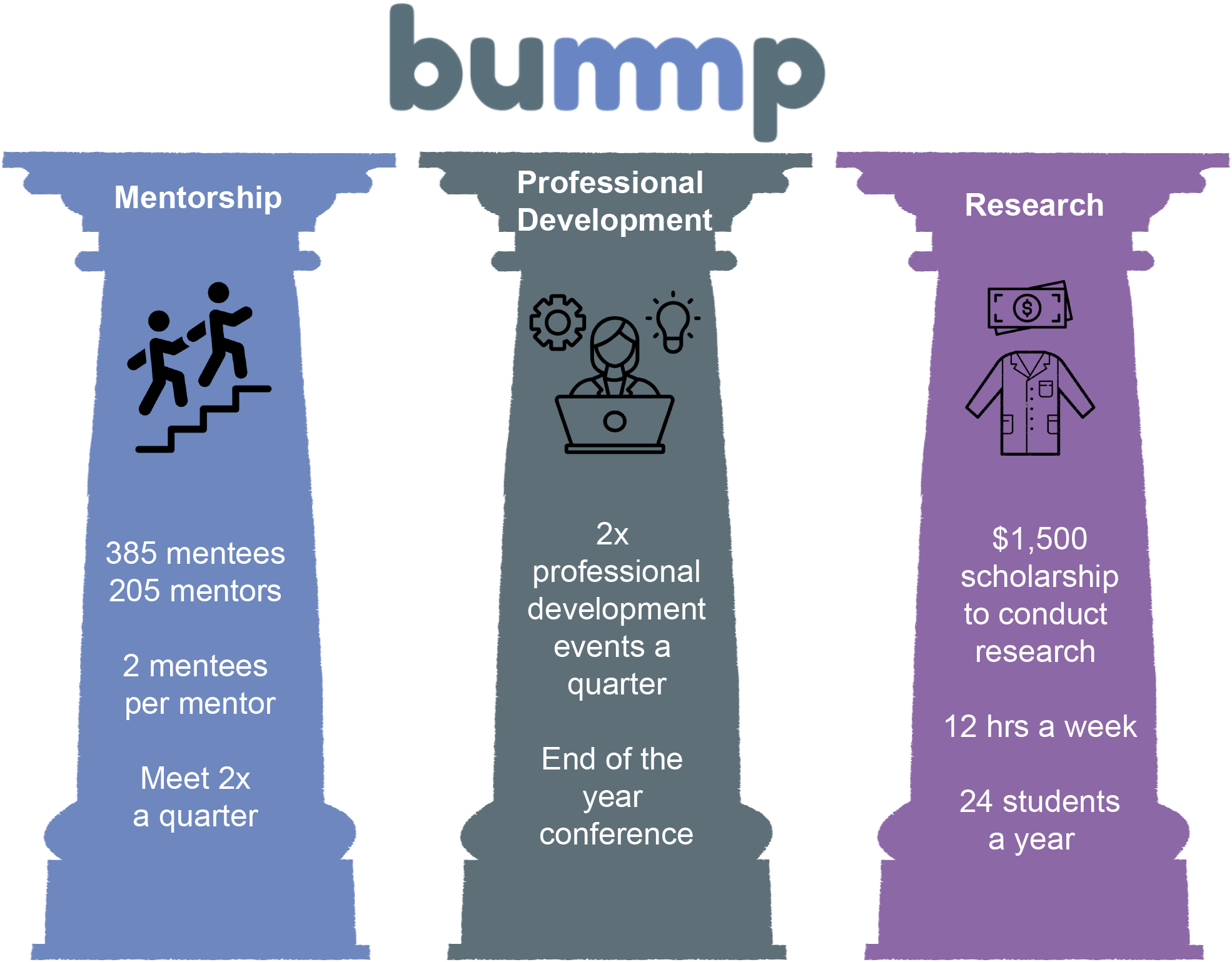
The Three Pillars of BUMMP. BUMMP is comprised of three main pillars - 1) Mentorship (blue), 2) Professional Development (gray), and 3) Research (purple). The three pillars provide different areas of support for our mentees.

## Materials & Methods

To capture both quantitative and qualitative data on the experience of mentees participating in BUMMP, surveys were utilized. The study obtained IRB approval and fully complied with ethical research standards.

Survey Design: Three surveys were utilized in this study.

Survey A: The Role of Mentorship Survey (BUMMP Mentees)

Survey B: The Role of Research Survey, BUMMP Research Apprenticeship Program (BRAP) (BUMMP Mentees who were or currently are BRAP recipients)

Survey C: The Role of Research Survey, Research Methodologies Training Lab (RMTL) (BUMMP Mentees who were RMTL participants in 2023)

The surveys aimed to assess students’ STEM persistence, confidence, acquired skills, and sense of belonging. Survey questions were selected from the Persistence in the Sciences Assessment (Hanauer et al., 2016; Hanauer & Hatfull, 2015), which includes questions about student belonging, identity, and efficacy. Additional questions relevant to mentees’ experience within the specific program (BUMMP, BRAP, or RMTL) were also included. The survey asks for self-reported demographics (race/ethnicity, gender, and parental educational background). Surveys were designed and distributed through Qualtrics.

### Participant recruitment

Students registered as BUMMP mentees in the 2022-2023 and 2023-2024 academic years were invited to participate in the study via email. The email included the research study’s purpose, participation expectations, and an exempt information sheet. When participants clicked the link to the survey, the survey introduction informed them again about their voluntary participation, and by completing the survey, they were giving their informed consent as research participants. A second participation invitation email was sent to current or former BRAP recipients to complete Survey B. A third participation invitation email was sent to mentees who participated in RMTL to complete Survey C.

### Respondents

Participants could skip any question in the survey, which accounts for the varying number of responses per question.

**Survey A:** Total respondents = 174

Academic Year (N = 169):

| 1<sup>st</sup> year | 2<sup>nd</sup> year | 3<sup>rd</sup> year | 4<sup>th</sup> year | 5<sup>th</sup> year | Master's |
| --- | --- | --- | --- | --- | --- |
| 33% | 14% | 24% | 21% | 1% | 7% |

Parental education background (N = 170):

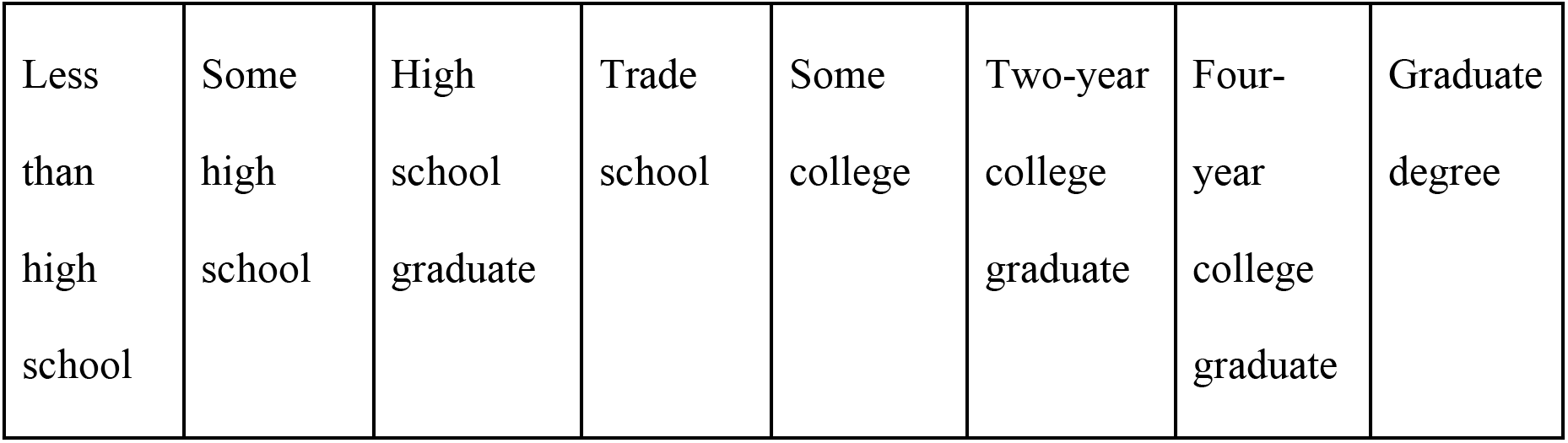
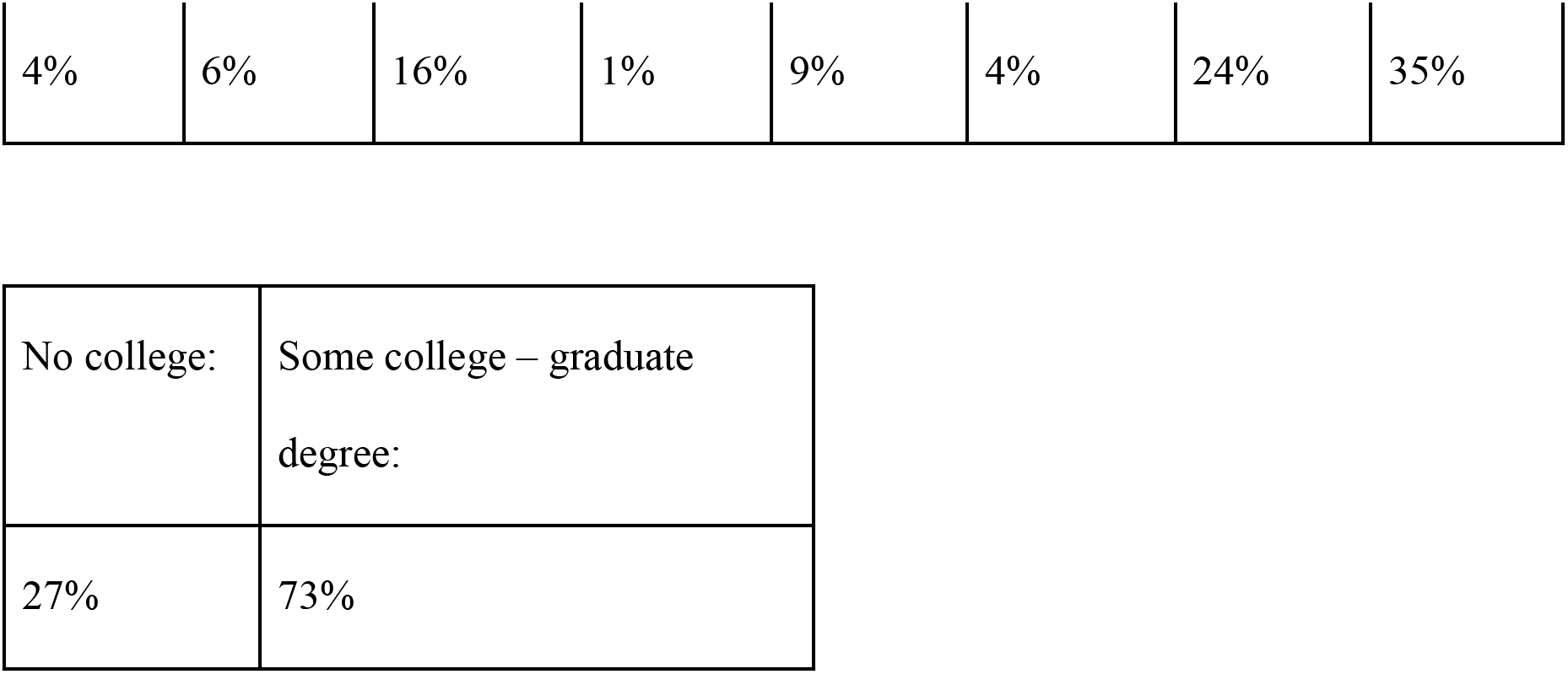

| Less than high school | Some high school | High school graduate | Trade school | Some college | Two-year college graduate | Four-year college graduate | Graduate degree |
| --- | --- | --- | --- | --- | --- | --- | --- |
| 4% | 6% | 16% | 1% | 9% | 4% | 24% | 35% |

|  |  |
| --- | --- |
| No college: | Some college – graduate degree: |
| 27% | 73% |

**Survey B:** Total respondents = 19

Academic Year (N = 14):

|  |  |  |  |  |  |
| --- | --- | --- | --- | --- | --- |
| 1<sup>st</sup> year | 2<sup>nd</sup> year | 3<sup>rd</sup> year | 4<sup>th</sup> year | 5<sup>th</sup> year | Master's |
| 7% | 7% | 0% | 64% | 0% | 21% |

Parental education background (N = 19):

|  |  |
| --- | --- |
| No college: | Some college – graduate degree: |
| 37% | 63% |

**Survey C:** Total respondents = 10 Academic Year (N = 8):

|  |  |  |  |  |  |
| --- | --- | --- | --- | --- | --- |
| 1<sup>st</sup> year | 2<sup>nd</sup> year | 3<sup>rd</sup> year | 4<sup>th</sup> year | 5<sup>th</sup> year | Master's |
| 0% | 25% | 38% | 13% | 13% | 13% |

Parental education background (N = 9).

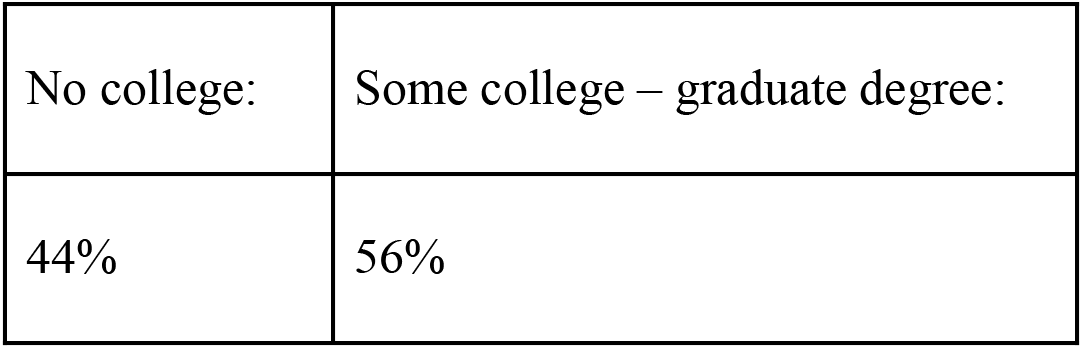

*Quantitative analysis:* Generalized linear models will be built to determine how student sense of belonging, self-efficacy, and STEM identity are functions of self-identified demographic information. All models will be assessed for overfit using AIC and ‘best’ models will be tested using restricted maximum likelihood.

## Results

### I. Mentorship

Studies have shown that mentorship helps students from historically excluded groups navigate academia by receiving thoughtful guidance and advice. Mentors who commit themselves to the success of their students and students who join mentoring programs have higher rates of success (Mariano et al., 2019; Marshall et al., 2022; Reeves et al., 2023; Rinderknecht et al., 2023). Students from historically excluded groups can often feel as though they do not belong in STEM spaces, which leads to a decrease in STEM retention through graduation. Dedicated mentorship alleviates this by bringing their mentees into STEM communities, guiding them through their undergraduate experiences, and reassuring their place in STEM. With this in mind, BUMMP aims to secure the success of UC San Diego’s historically excluded students by emphasizing mentorship as an integral component of our program.

#### Recruitment of mentors

Recruitment of high-quality mentors is necessary to maximize the resource network of our mentees. To this end, BUMMP recruits mentors directly from the rich scientific community at UC San Diego, pairing undergraduate and master’s students with a mentor, who is typically a graduate student, post-doctoral fellow or faculty member at UC San Diego. Most recently, industry scientists from San Diego have become valuable mentors. Our strong pool of mentors allows our mentees to develop a relationship with someone who has experience in STEM and thus, can offer them sound career advice, guidance, and connections to strong networks within the STEM community in the United States.

At the start of the academic year, both mentees and mentors are recruited to join BUMMP. Our recruitment information is distributed to professors and instructional assistants, undergraduate newsletters, and coordinators of departmental listservs for all biology-related fields (biological sciences, biomedical sciences, bioengineering, biochemistry, bioinformatics, and the MD/PhD program). Additionally, we strategically recruit mentees from historically underserved communities by enlisting the assistance of the UC San Diego Resource Centers which are designated to service an underserved group. We use online surveys (google forms, qualtrics, survey monkey) to gather interest in the program, and all the interested parties are invited to attend an informational session. To officially become a BUMMP mentor or mentee, everyone is then required to submit their scientific interests/goals and contact information through our Intake Survey. To maximize the impact of our program, there is no GPA or experience requirement to join BUMMP. The singular requirement is that each BUMMP mentee and mentor must attend mandatory training sessions. These training sessions are strategically designed to ensure our mentors and mentees are up to date on professional standards and principles of culturally responsive mentorship, which will allow them to be better mentors (Mariano et al., 2019; Marshall et al., 2022; Romney & Grosovsky, 2023). Having completed our intake survey and attended these training sessions, members are now eligible for pairing. We have found this approach to be effective in increasing attendance at events and ensuring that pairings remain active throughout the year.

#### Mentor-Mentee pairing

To customize the mentorship experience, we use an algorithm that automatically matches mentors and mentees first by gender preference, and then by primary area of scientific interest. Secondary interests are used for mentees who cannot be successfully matched due to insufficient numbers of mentors in their primary interest. Generally, each mentor is assigned to two mentees. Once the maximum number of pairs have been made, a public spreadsheet is shared with all mentees and mentors. The mentors are responsible for using this information to reach out to their mentees and set up the first mentoring session. We encourage mentors to initiate meetings with their mentees, as reaching out to mentors can be intimidating for our mentees and shows the investment of the mentor to the mentee. This past year we had 385 mentees and 191 mentors who joined our BUMMP community.

#### Mentor-Mentee pair tracking

Each mentee-mentor pair is required to meet twice a quarter to uphold the consistency of mentorship. Twice a quarter, mentees fill out a survey describing the outcome of their meetings. At the end of each quarter, mentors fill out a survey describing their meetings and pairings. The purpose of this is two-fold: 1) it increases participation, ensuring the meetings are being held, and 2) it allows us to get feedback on how mentors and mentees perceive the program and their pairing. This feedback is reviewed both quarterly and at the end of each academic year so that necessary suggestions can be incorporated into the program design. Our program coordinator monitors and tracks our mentor-mentee meetings to ensure that our members are consistently meeting, and that our mentees are getting the mentorship they need. Additionally, mandating attendance for our mentors and mentees at our mandatory training sessions ensures they understand their duties and fosters accountability from the outset of the year.

#### Community building and expanding our mentorship network

In addition to strong mentorship, HEG students benefit from stable supportive communities of peers with similar experiences. Therefore, we have intentionally created a safe community where mentees can meaningfully engage with each other, and with their mentors. We use Slack as a platform to encourage mentees to ask questions about ideas included but not limited to STEM, career advice, scholarship opportunities, and graduate or professional school. To strengthen the BUMMP community, we have quarterly general body meetings and social gatherings are held where all members can come together, often sharing meals, and partaking in recreational activities. Moreover, to encourage closer relationships with additional members of BUMMP outside of the mentor-mentee pair, we created “pods” where 3 sets of pairings are grouped together, usually by research interest. This approach offers a bonus benefit by giving mentees access to a wider network of mentors. Additionally, we have BUMMP-branded items that we give to our members free of cost, including shirts, stickers, folders, and tote bags.

#### Mentorship outcomes

To determine the effectiveness and impact of mentorship on BUMMP mentees, a survey was distributed to each student in the mentorship program for the Fall and Winter quarter of the 2023-2024 academic year. In the mentorship program, productivity is reflected in mentee perception of the program, specifically mentee satisfaction, impact of the mentor, and the fulfillment of their expectations. Highly productive mentor pairs are those who have a score of at least 7 out of 10 for these metrics. 87%, 74%, and 83% of the respondents reported that their mentor pairings were highly productive in these three areas, respectively (Figure 2A). Supporting these data are testimonies given by our students (Table 1). Overall, these findings reassure us that the mentorship program is enriching to the mentees. Further, we ascertained the specific areas of productivity that the mentorship program addressed. The areas mentees felt they received the most support were in career planning (37%), academic planning and preparation (19%), and research (~16%) (Figure 2B). In addition to guiding the content and structure of the BUMMP mentorship training, this data suggests that BUMMP mentors, who have years of lived experience working in STEM, are already highly qualified to provide mentorship in these areas, leading to mentee satisfaction with their pairings. Moreover, this satisfaction was equally enjoyed by first-generation and non-first-generation students, suggesting that quality mentorship benefits students of differing backgrounds (Figure 2C). This is critical as the mentorship program serves students of varying parental education backgrounds (Figure S1A). Further, participation in the mentorship program has resulted in 35% (50 out of 145) mentees receiving an opportunity, such as a company internship or research position in a UC San Diego lab, despite the fact that 70% of these respondents have only been in the mentorship program for two quarters (Figure 2D, Figure S1C). This highlights the power of networking and pristine mentorship, even in the short-term, to embolden mentees to take important steps in their STEM career. Our mentees received support from various areas from their mentors. Taken together, this data supports the adeptness of our mentorship pairing algorithm to select the most productive mentor-mentee pairs.

**Figure 2:**
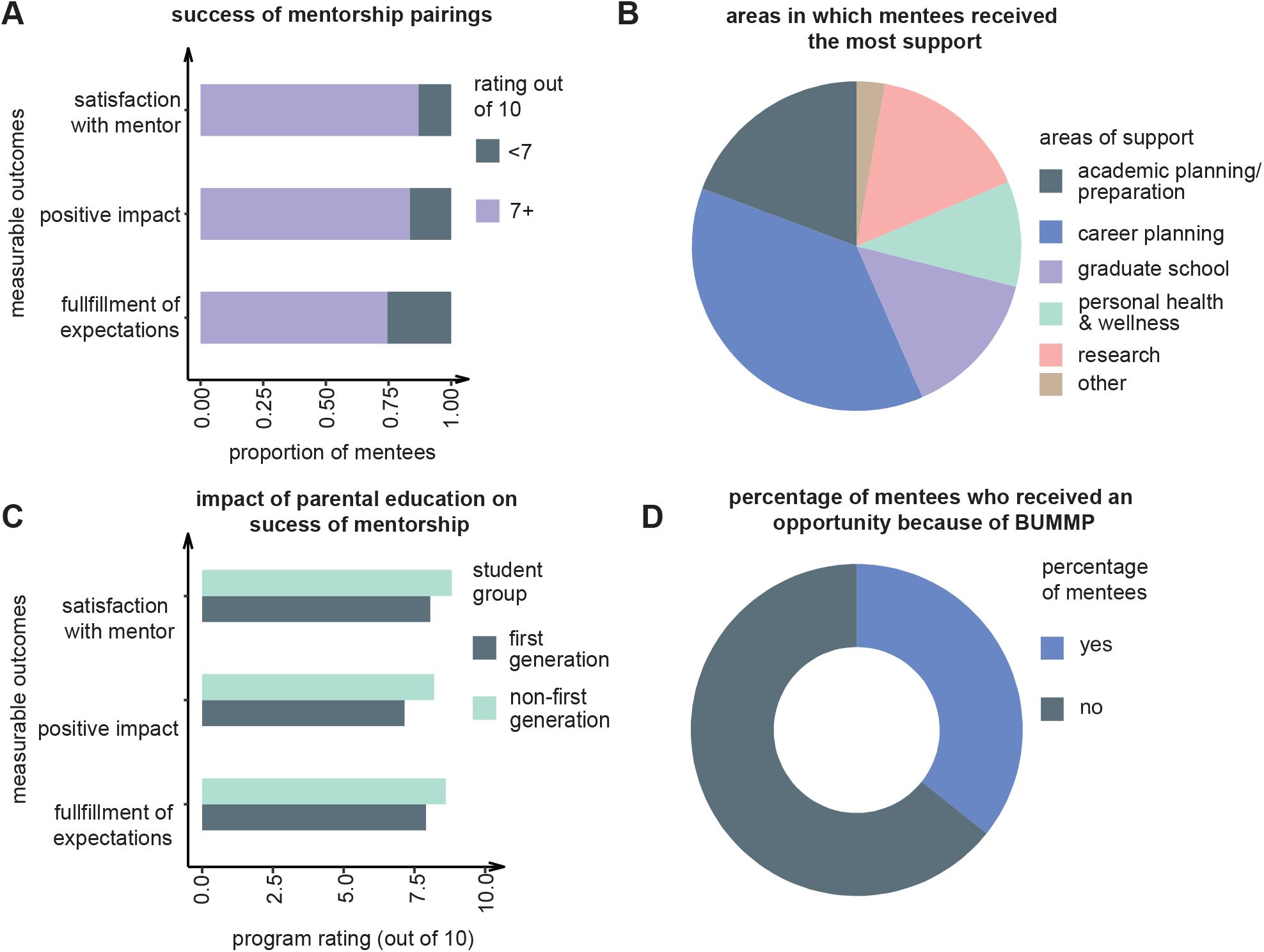
BUMMP mentors have a positive impact on mentees in the mentorship program. (A) Opinions of mentees in the mentorship program on the success of their mentorship pairing. Mentee self-ranking on a scale of 1-10, 10 being the highest. (B) Breakdown of the areas of support mentors assisted mentees within the mentorship program. (C) Opinions of mentees in the mentorship program, who are first generation (gray) or non-first generation (green), on the success of their mentorship program. Mentee self-ranking on a scale of 1-10, 10 being the highest. (D) Percentage of mentees in the mentorship program who received a STEM-related opportunity because of being in BUMMP during the first two quarters of the 2023-2024 academic year.

**Table 1:** Testimonials from BUMMP Mentees.

|  | Mentee testimony |
| --- | --- |
| Mentorship | “BUMMP has provided me with so many resources and invaluable guidance, not to mention the passionate and caring mentors!”<br>“I am super excited to keep on working with my mentor and am extremely thankful for the opportunity I have of getting to have a mentor I can talk to for guidance.” |
| Events | “I am so grateful for all the advice I received on both how to succeed in interviews and how to build experience as a first-year undergraduate student. Mentors were so willing to provide me with tips from their own experiences in labs and working with undergraduates.”<br>“I like the chance of gaining tips and confidence from professors. I think that words of affirmation are important and it is important for students to recognize that professors are just approachable and supportive mentors.” |
| BRAP | “I have really enjoyed my time as a BRAP scholar and feel that BUMMP in general is one of the only places throughout my academic career thus far where I have felt fully supported. I have struggled a lot with imposter syndrome/do not get a ton of support for what I do, and BUMMP has been a nice reminder that there are people in my corner. Being a part of BRAP has solidified that support for me, as it has recognized all of the effort and time I have put in.”<br>“My time as a BRAP scholar really inspired me to further a career in research. Having both BUMMP and BRAP supporting your research during this time gave me the resources to financially support myself and the community to share my excitement for research with.” |
| BUMMP | “BUMMP helped me get the most out of my experience at UCSD. I have been able to find a lab to volunteer in, and I will be learning some basic techniques I need to know and understand. I have also received help when it comes to applying to graduate schools.”<br>“BUMMP has been an amazing program for me and has helped me gain confidence and build a support network of people all seeking to further scientific and medical advancement which is so so cool. So thank you! |

Science identity is an excellent predictor of student retention in STEM, especially students from HEGs (Hanauer et al., 2016; Hanauer & Hatfull, 2015). We observed that multiple aspects of science identity were positively impacted by participation in the mentorship program, including mentees’ sense of belonging to the community of scientists (80%), feeling that their identity and background are welcomed in their academic program (88%), feeling like they belong in science (90%), their perception of the daily work of scientists (83%), sense of value as a scientist because of their identity and background (71%), and certainty of their place in science (68%). (Figure 3A). These data suggest that having a knowledgeable mentor in STEM and being part of a safe community of scientists in a program like BUMMP has a significant positive impact on their science identity. Further, both first generation and non-first-generation students experienced positive impacts on their science identity, suggesting that strong mentorship and community can help make all students feel welcome in STEM (Figure 3B).

**Figure 3:**
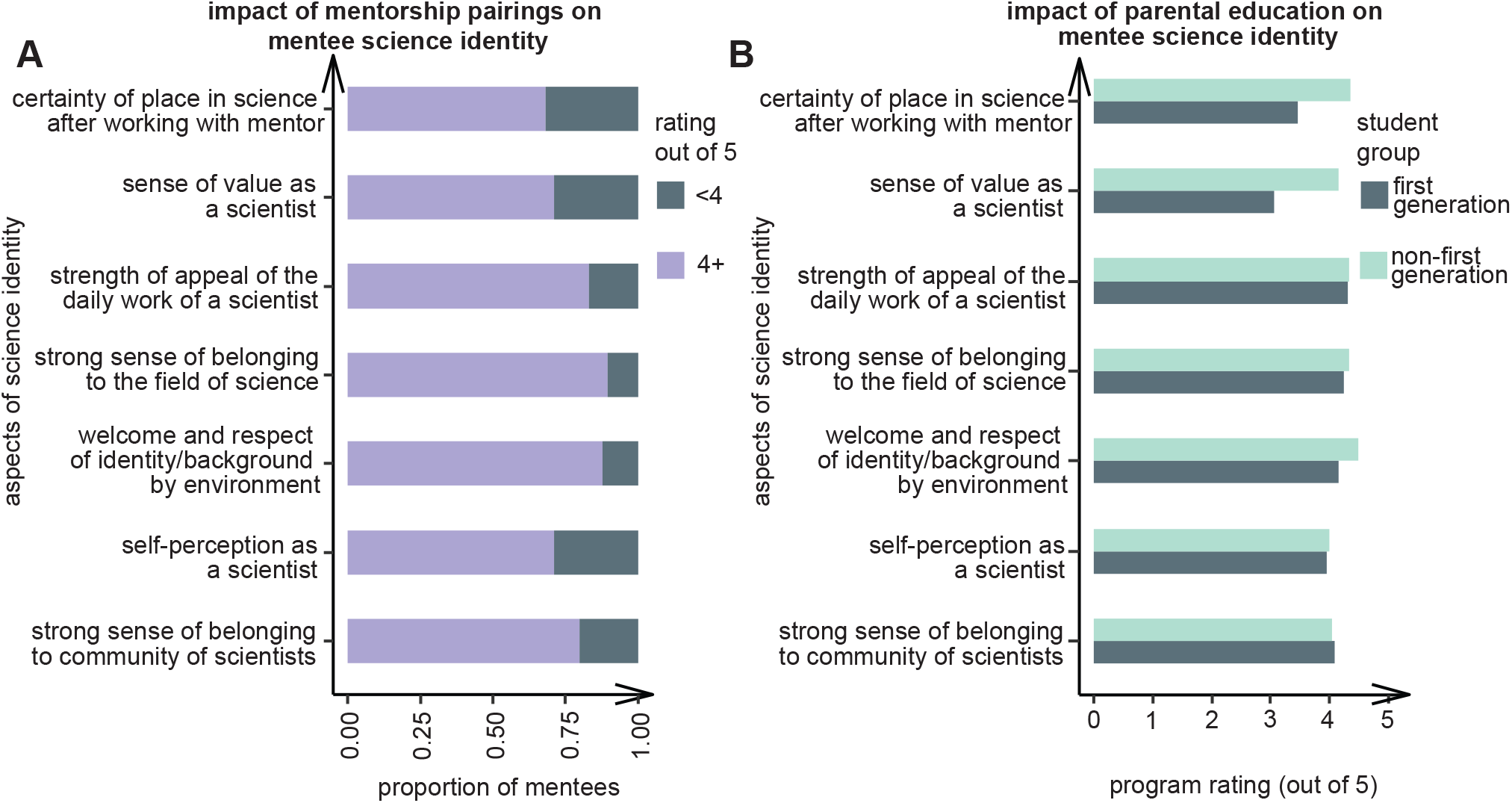
BUMMP mentors positively impact mentee science identity. (A) Impact of mentorship pairings on mentee science identity. Mentee self-ranking on a scale of 1 to 5, 5 being the highest. (B) Impact of mentorship pairings on mentee science identity for first generation (gray) or non-first generation (green) mentees. Mentee self-ranking on a scale of 1 to 5, 5 being the highest.

Finally, the mentorship program is equally beneficial for the mentors. In the mentor testimonials, mentors describe their following experience: “I feel more connected to the UCSD Biology community by being a mentor. It’s been incredibly rewarding to watch [their] mentees grow and achieve their goals and help facilitate that for them”, and “I really like that I get to talk and connect with undergrads on campus. I think it is very rewarding to be able to help mentees and help them figure out how to get started in science and just make a connection with them. I think BUMMP is very valuable in facilitating those connections.” (Table 2). BUMMP provides a welcoming community that not only benefits our mentees, but our mentors as well. Many of our mentors are graduate students, and this allows them to get to know other scientists and faculty at UCSD. Further, it fosters connections at all levels. Many of our mentors know the value of mentorship because it helped them in their own careers, and BUMMP provides them with an avenue to give back to students in a meaningful way.

**Table 2:** Testimonials from BUMMP Mentors.

|  | Mentor testimony |
| --- | --- |
| Mentorship | “Mentors were influential and a necessity in helping me achieve my goals and further my scientific career. I could not have gotten this far without them, and BUMMP allows me to support others in the same way.”<br>“I really like that I get to talk and connect with undergrads on campus. I think it is very rewarding to be able to help mentees and help them figure out how to get started in science and just make a connection with them. I think BUMMP is very valuable in facilitating those connections.” |
| Events | “Loved the mentor training! Thought it was very informative.”<br>“The power point presentation during the interview workshop was insightful and comprehensive. The group sessions were fun and useful. As a mentor it was pleasant to learn about different students background and research experience.” |
| BRAP | “When I was a graduate student, an undergraduate I mentored in lab received the BRAP award. This award allowed her to continue to develop as a scientist during the academic quarter while being financially recognized for her effort and dedication in lab. This student has now gone on to receive a masters in biology and plans to pursue an MD-PhD; both opportunities which have been made possible by the experience she gained as a BRAP recipient.”<br>“As a new PI, the first BS/MS student in my lab was awarded BRAP scholarship. The support from BRAP allowed the student to focus on his research and eventually complete his master's degree in my lab. The student is currently a Research Assistant in my lab and will be starting medical school in the fall. BRAP was instrumental to allowing him to achieve these goals.” |
| BUMMP | “I think that the mission of BUMMP is great and desperately needed in academia. BUMMP matches many students with research labs and funds amazing undergraduate research. I think BASS was great too.”<br>“I feel more connected to the UCSD Biology community by being a mentor. It's been incredibly rewarding to watch my mentees grow and achieve their goals and help facilitate that for them.” |

### II. Professional Development

Both within and beyond academia, the development of professional and career skills is linked to career success and retention (Blau 2017; Robinson et al. 2016). Such skills comprise career knowledge, time management, leadership, problem solving, decision making, and various interpersonal skills. Functionally, there is a clear need for professional skills in both academic activities and career planning. Communicating with professors and classmates, collaborating with peers, and involvement in student organizations can become more challenging without a developed professional skill set. After graduation, these skills help in understanding career and postgraduate options, submitting applications and interviews, and realizing career success. For now, many of these soft skills are not formally incorporated into undergraduate curricula. Data collected by the Black Academic Excellence Initiative Committee at UCSD shows that a lack of development opportunities on campus presents challenges to members of the Black undergraduate community (Berthoud et al. 2021). Even outside of a university setting, historically underserved students do not have equitable access to professional development opportunities. Poverty, underfunded high schools, family instability, and lack of mentorship are all contributing factors to professional uncertainty (Lindstrom et al. 2022). As a result, underserved students and their teachers consider limited availability of professional development resources to be a barrier to career readiness (Lindstrom et al. 2022).

BUMMP aims to address the gaps that persist in career readiness and scientific identity for historically underserved students. As an organization, we foster a diverse community which serves two roles: (1) to establish a network of support that offers representation and acceptance in STEM and (2) to help students develop professional and career skills through community-led workshops. To accomplish this, BUMMP organizes multiple events per academic quarter for the BUMMP community, which is described below.

#### Annual Mentor and Mentee Training

Cultivating a community where underserved students feel safe and respected is a priority of BUMMP. That is why all mentors and mentees are required to complete our mentor or mentee training prior to being paired. It is important to us that our mentors understand the barriers underserved students face, as well as the diverse backgrounds they come from. For this reason, our mentor training is designed to educate mentors on these topics and provide tangible guidance on effectively mentoring students from different backgrounds. In our training, we introduce mentors to culturally responsive mentorship, in which mentors embrace and learn about the cultural and social identities of their mentees. This method of mentorship is shown to be mutually beneficial for mentors and mentees. For mentees, acceptance of non-STEM identities helped students feel more confident as researchers and become more committed to their degrees (Haeger & Fresquez, 2016). This is reflected in our own program, in which many mentees agreed that their mentor valued and respected their identity outside of science (Figure 3A). Meanwhile, mentors report feeling more confident in having conversations about race and culture with their students and experience improved reciprocal relationship building(Haeger & Fresquez, 2016; Pfund et al., 2022). Importantly, mentors are walked through the principles and benefits of culturally responsive mentorship and given guidance on initiating and responding to conversations on race and culture with their mentee. Meanwhile, mentees undergo training which introduces the concept of culturally responsive mentorship and information on how to get the most out of mentorship and BUMMP. This involves guidance on communication practices with mentors, expectations for both mentees and mentors, and BUMMP and campus resources. These trainings allow us to build a community with a greater understanding and acceptance of different backgrounds and lays a foundation to set our mentees and mentors up for success.

#### Professional Development Workshops

With persisting gaps in access to professional development resources for underserved students (Berthoud et al. 2021; Lindstrom et al. 2022), BUMMP makes it a priority to provide students with opportunities to build professional and career skills. Notably, we hold several workshops a year including writing/application, interview, and career workshops. Each of these is tailored to provide foundational information on application preparation, interview practices, and career options respectively. More importantly, BUMMP mentors are invited to lead small groups during these workshops, providing students with invaluable first-hand experience and one-on-one feedback on their work. In addition to providing access to professional and career skills, this allows mentees to expand their network of support and connect with other mentors and mentees in the program. As a result, we have received glowing testimonials from both mentors and mentees (Table 1, Table 2).

#### Research Blitz

In addition to mentorship, research experience helps students build science identity and improves retention in STEM (Estrada et al., 2018). However, understanding how to join a research lab can be unintuitive and overwhelming – particularly for students struggling with science identity. Meanwhile, many labs greatly benefit from undergraduate researchers, but similarly struggle to find students. To remedy this, BUMMP hosts a ‘Research Blitz’ where faculty members that want to mentor undergraduate students are invited to present a poster. Students interested in joining labs or learning about research on campus are encouraged to attend and connect with prospective labs. This gives students the opportunity to narrow down research areas of interest and network with faculty, post-docs, and graduate students. In addition, BUMMP has created our own poster detailing steps to take to find research positions on campus. In the past, these events have facilitated helping many students get their first research position.

#### BUMMP Annual Student Symposium

Finally, BUMMP organizes the BUMMP Annual Student Symposium (BASS) at the end of the academic year. This conference is the culmination of mentee and mentor achievements and BUMMP’s core values. Throughout the day, mentees are given the opportunity to give a research talk or present in a judged poster session. This allows mentees to showcase their accomplishments in a supportive environment and practice scientific communication skills. Additionally, we include an Opportunity Fair where we invite graduate programs, internships, student organizations, and summer research programs to promote their programs. This provides students exposure to resources on and off campus and gives opportunity for face-to-face networking with other organizations. Finally, the conference concludes with presenting awards for the best poster presentations and merit awards. This event is the ultimate celebration of BUMMP, our accomplished students, and our dedicated mentors.

#### Professional Development Outcomes

Barriers unique to HEGs can often contribute to feelings of inadequacy and deeply impact student’s sense of belonging, and therefore retention, in STEM. This is exacerbated by negative social influence and systemic oppression (Estrada et al., 2018; Tello & Goode, 2023), and can culminate into feelings of not belonging in STEM. Meanwhile, studies have shown that underserved students that develop a scientific identity (i.e. feelings of belonging in a scientific community) are more likely to remain in a STEM career beyond graduation (Estrada et al., 2018). During participation in BUMMP, 80% of students agreed that they had a strong sense of belonging to the community of scientists. In turn, 71% agreed they had come to think of themselves as a ‘scientist’ (Figure 3A). In this way, our commitment to community supports students in the development of their professional skills and science identity in order to support their retention in STEM beyond their undergraduate education.

### III. Research

Undergraduate research is key in shaping students’ trajectories within STEM fields, offering not only academic enrichment but also fostering a diverse range of skills essential for their professional growth. Participation in undergraduate research not only enhances students’ retention in STEM majors and interest in pursuing graduate degrees but also fosters crucial skills essential for their professional development. Participating in research increases students’ retention in a STEM major and their interest in pursuing a graduate degree (Hathaway Biren, A Nagda Sandra R Gregerman 2002; Lopatto 2007). Participation in research has been shown to have a positive impact on graduation rates for students coming from underserved backgrounds (Haeger et al., 2024; Haeger & Fresquez, 2016). Additionally, the vast majority of students participating in undergraduate research express interest in pursuing a graduate degree in science (Lopatto 2007). Students participating in research indicated they felt their experience better prepared them for a career in STEM because they better understood the research process and gained skills relevant for their career path such as data analysis, lab techniques, and reading primary literature (Lopatto, 2007). Participation in research is not only useful for building up skills that are relevant to a career in STEM, but it also important for building up general professional skills such as independent thinking, problem solving, curiosity and presenting (Russell et al. 2007; Thiry et al. 2011).By actively engaging in research endeavors during their undergraduate years, students not only enhance their retention in STEM majors and bolster their motivation towards pursuing advanced degrees but also acquire a multifaceted skill set encompassing technical skills, critical thinking, problem-solving, and effective communication, thus priming them for success in both academic and professional realms.

#### Inequalities in access to research opportunities

Access to research opportunities in STEM fields remains unequal. Key barriers that students from HEGs in STEM face are lack of role models, financial constraints, and cultural barriers (Pierszalowski et al., 2021). These barriers are especially pronounced for first-generation students in college, students coming from a low-income background, and students identifying with underrepresented groups (Stebleton and Soria 2016; Kim 2009). Identifying these opportunities is often a difficult task for undergraduate students, especially those from underrepresented backgrounds. Research has shown that some of the main difficulties for students looking for research opportunities is that there are limited centralized resources that advertise such opportunities, and a limited number of opportunities to begin with (Pierszalowski et al., 2021). Addressing these systematic disparities is important for fostering inclusivity and leveling the playing field, ensuring that all aspiring scientists have equitable access to the experiences and opportunities that research involvement affords.

To provide financial support for our students, we have created multiple scholarship, fellowship, and research opportunities. One of BUMMP’s main research scholarship programs is known as the BUMMP Research Apprenticeship Program (BRAP). BRAP was created to support low-income, first-generation college students, and students from groups underrepresented in graduate education that are interested in participating in research at UCSD by providing quarterly research stipends and educational programming. For students looking to gain practical skills that can be applied to both academic and industry lab settings, and who are unable to commit to a research experience, we have also actively partnered with the Research Methodologies Training Lab (RMTL) which is led by the UCSD School of Pharmacy. This program allows students to gain a variety of technical skills during a ten-week course. Finally, to continue to mitigate financial burdens faced by our students, we also offer a yearly merit-based award.

#### The BUMMP Research Apprenticeship Program

Participation in research has been shown to improve URM retention in science as it contributes to the students’ development of their scientific identity. However, many undergraduate research positions are volunteer positions, or provide course credit as compensation. According to a 2016 survey from UCSD, 32% of student researchers indicated that their research position was volunteer based and another 54% indicated that their research was for course credit. Only 14% of students participating in research were paid (UCUES survey 2016). This practice of not providing monetary compensation for undergraduate research makes participating in research a hurdle for those who have other financial obligations, such as those coming from a low-income background. BRAP is a scholarship opportunity offered by BUMMP, which allows students to devote more time to research opportunities that they would have otherwise needed to devote to other jobs that provided a stable source of income. It prioritizes funding to low-income, first-generation college students, and students from groups historically underserved in graduate education that are interested in pursuing research during the academic year. This scholarship awards up to $1,500 per quarter to help expose students to cutting-edge research in labs under the guidance of a principal investigator. In addition to providing scholarship opportunities, students are to provide a written report or oral presentation at the end of their research experience. To facilitate this in welcoming and supportive environments, students are invited to present their work at BUMMP’s general body meetings (GBMs) as well as the BUMMP annual student symposium.

To develop a scholarship program, support is needed from many different stakeholders (see “Keys to Success” section for more details). BUMMP has a prolonged scholarship program in which students apply during the first quarter of the year, and if awarded they obtain funding for the entire year (3 quarters). Most students participating in research participate for multiple quarters, so this ensures that they have continued financial support for their research opportunities. Students also have the opportunity to apply during the second quarter of the year, and obtain funding for the duration of the year. Depending on the funding mechanism that a program uses, it will always be useful to identify other support systems for students, such as other funding systems and internship programs as this provides students with multiple opportunities to achieve the finances they need to pursue research opportunities.

#### Research Methodologies Training Program

BUMMP has partnered with the Research Methodology Training Laboratory (RMTL) program at UCSD to provide students with formal lab research experience in an educational setting. Through this program, first-generation students and students from historically underserved backgrounds who lack formal lab research experience gain training in skill sets that are fundamental to work in industry and academia. RMTL is a ten-week, fully funded on-campus fellowship that students participate in during the academic year. Throughout the course of the program, students gain 80 hours of basic science training. Additionally, they gain extra mentorship from the program staff, are invited to workshops specific to the program, and participate in networking events separate from those hosted by BUMMP. At the end of the fellowship, students earn a lab assistant certificate and have the opportunity to give a poster presentation.

#### Application and Review Process

Our application questions are geared at understanding a student’s background and aspirations, rather than their success thus far. Students are asked to write a statement of interest and a statement of obstacles. Through the student’s statement of interest and statement of obstacles, students communicate how BUMMP and the scholarship will help them achieve their goals and what obstacles they have overcome so far to reach their current point in their career. In addition to those two main application questions, students also write a project blurb and must submit a document of faculty support. These two documents are the primary way for reviewers to assess whether the student is actively participating in a non-funded research project. The BRAP scholarship is only offered to students who are actively involved in BUMMP’s mentorship program. To help students identify research opportunities, we host events geared at allowing students to learn more about the research opportunities on campus, in the surrounding industry, and at surrounding research institutions. Further, the program also shares opportunities that our routed to us through email and social media.

When reviewing applicants, BUMMP takes a holistic approach. Students that self-indicate that they are first-generation college students or come from a low-income background are prioritized. This is a priority for BUMMP as previous research has shown that these groups are underrepresented in STEM and higher education. Additionally, to ensure that BRAP and RMTL have a wide reach across students participating in BUMMP, our scoring process prioritizes students that have not previously been awarded. This maximizes the impact of BUMMP across students and ensures that we are always providing opportunities to students who have not yet had the opportunity to gain key research experiences. In line with this, BUMMP prioritizes students in later years of their undergraduate career to participate in the RMTL program as they will soon be applying to academic, or industry jobs and the hands-on training obtained during the RMTL program can be marketed as important skills during their application process.

#### Research Outcomes

Students who participated in BRAP and RMTL come from diverse backgrounds. 66% of participants in BRAP and 55% of RMTL participants indicated that they were first generation college students (Figure S1B, Figure S2&B).

Participation in BRAP overall had a positive impact on participant’s ownership over their research project, their self efficacy, and their science identity (Figure 4). For example, 100% of students participating in BRAP felt that they had responsibility over a research project, and 93% felt that participating in BRAP positively impacted their ability to overcome challenges during a research project (Figure 4A). Regarding self-efficacy, 100% of students participating in BRAP felt confident in their ability to form hypotheses, collect data, interpret data, and in their technical skills (Figure 4B). BRAP also had a positive impact on a student’s perception of their science identity: 93% of BRAP participants indicated that they had a strong sense of belonging to the field of science and overall felt welcome and respected, and like they belonged within the community of scientists (Figure 4C). Overall, participation in BRAP had a positive impact on student’s career trajectories (Figure 4D). Students who participated in RMTL also had a similarly positive experience. The majority of students that participated in RMTL felt positively about their self-efficacy in science and their science identity after their participating in the program (Figure S2C&D). Mentees describe their following experience in BRAP: “I have struggled a lot with imposter syndrome/do not get a ton of support for what I do. … Being a part of BRAP has solidified that support for me, as it has recognized all of the effort and time I have put in. “My time as a BRAP scholar really inspired me to further a career in research. Having both BUMMP and BRAP supporting your research during this time gave me the resources to financially support myself and the community to share my excitement for research with (Table 1).”

**Figure 4:**
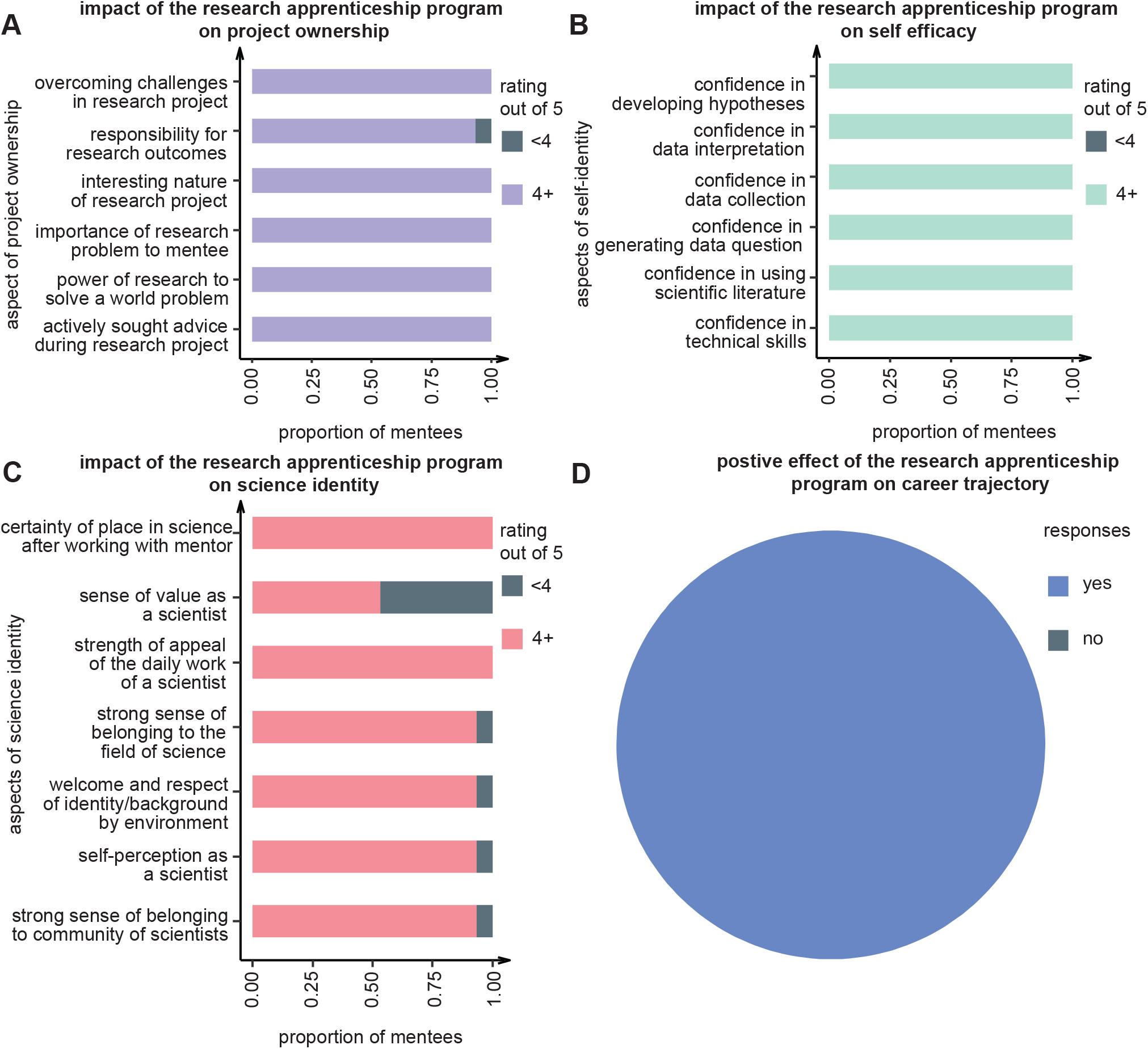
The BUMMP Research Apprenticeship Program (BRAP) has a positive impact on mentee scientific development and science identity. (A) Impact of BRAP on mentees project ownership. Mentees self-ranking on a scale of 1 to 5, 5 being the highest. (B) Impact of BRAP on mentee’s self-efficacy. Mentees self-ranking on a scale of 1 to 5, 5 being the highest. (C) Impact of BRAP on mentee’s science identity. Mentees self-ranking on a scale of 1 to 5, 5 being the highest. (D) Proportion of mentees in BRAP who think BUMMP had a positive effect on their career trajectory.

**Figure 5:**
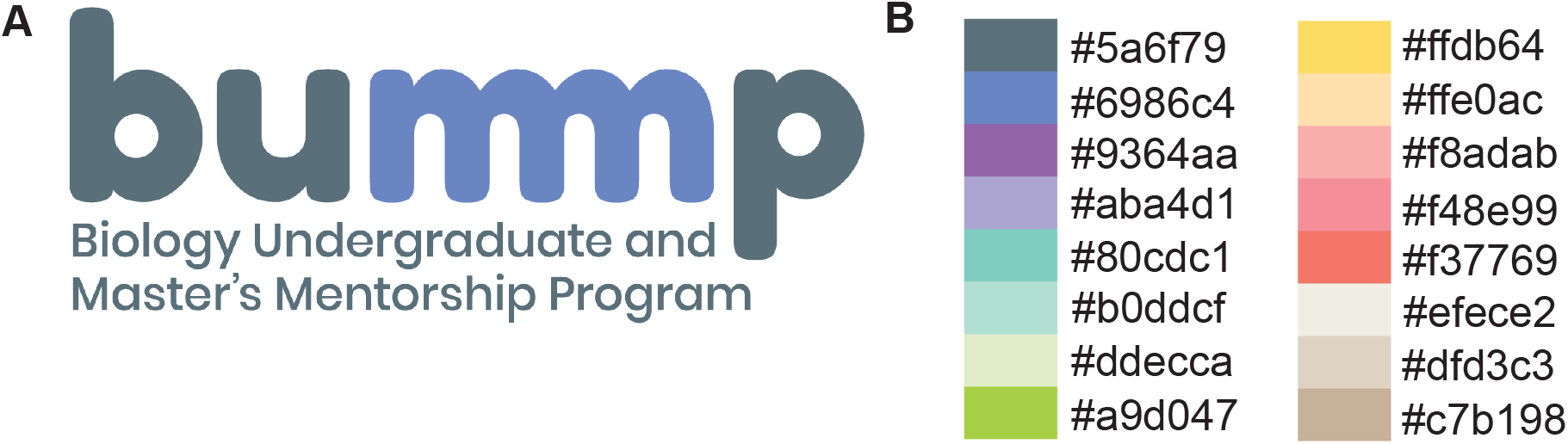
The BUMMP brand is strategically designed to foster a sense of community. (A) The primary BUMMP logo features interconnected m’s to symbolize boundless connections among mentors and mentees, and between mentees and opportunities. (B) A pastel color palette embodies the approachability and reliability of BUMMP.

### IV. Keys to success

BUMMP relies on several factors to ensure the effectiveness and sustainability of the program. The following were considered most important for program success:

#### 1. Commitment to BUMMP’s core values

The BUMMP leadership team is diverse, consisting of undergraduate students, graduate students, and a faculty member, who experienced a lack of sense of belonging and science identity in academia. Having experienced the positive impact of mentors on their own academic and professional development, the leadership team recognizes the pivotal role mentors play in helping individuals navigate the complexities of academia. This empathy becomes a driving force behind their commitment to supporting students. The shared lived experiences of the leadership team create cohesion, aligning everyone with BUMMP’s goals and objectives.

#### 2. A visible brand rooted in BUMMP’s core values

The BUMMP brand was intentionally designed to recapitulate our approach to increase the retention of HEGs in STEM, particularly our prioritization of the sense of belonging, safety, and community within every mentee. We curated a palette of soothing, pastel colors which are (1) muted to symbolize approachability and create a sense of belonging, (2) variable to represent all the brilliant diverse minds that gather from all walks of life in BUMMP, and (3) sufficiently vibrant to promote anticipation and excitement. Our logo concept displays two m’s joined together to represent the strong connections among our mentees and mentors, and endless opportunities. A soft blue is BUMMP’s main color and is a symbol of our dependability, authority, security, and trust.

#### 3. Sufficient and sustained support

To develop a scholarship program, support is needed from many different stakeholders. Key steps for starting this scholarship program include: identifying funding source, setting up an application and rubric, helping students identify research opportunities, and tracking involvement during the program. To go about obtaining funding, we partnered with the UCSD Biological Sciences Development Team. Development teams are essential and prevalent at many universities as they play a core role in obtaining financial support from university partners and alumni. In addition to pursuing funding through philanthropic individuals and organizations, foundations that are mission aligned with the program are also key funding sources. For BUMMP, this includes the Intuitive Foundation. Finally, the importance of increasing diversity in STEM is broadly recognized as important, and because of this many research funding sources add supplementary funding for diversity initiatives. We have been able to acquire funding from The Packard Foundation, The Chan Zuckerberg Initiative, and the NSF through supplementary grants from supportive principal investigators at UCSD. In addition to identifying opportunities for funding, it is also important to work with your institution to ensure timely receipt of finances. For example, there could be multiple ways that a funder may be able to give to the organization through the school, and those details must be figured out prior to funding. Over the past 4 years BUMMP received: $180,000 of donor funding, $105,000 of private funding, $30,000 of the institution’s internal grants, and $120,000 of non-profit organization funding. Moreover, the institution recruited a full-time program staff (see below) by year 3. These activities emphasize that inclusive excellence is an institutional priority.

#### 4. Having a full-time program coordinator

The BUMMP program is run by graduate students who are not able to dedicate themselves full-time to daily operational tasks. To ensure longevity, it is critical to have a program coordinator. A program coordinator oversees research and merit scholarships. They play a crucial role in ensuring the smooth administration, distribution, and effective utilization of scholarships within the program, contributing to its overall success and impact. Students greatly benefited from the guidance of a dedicated staff coordinator who addressed inquiries regarding eligibility and the scholarship application process. The coordinator efficiently manages reimbursements for daily BUMMP activities and maintains a direct connection with upper administration and leadership across different departments and campuses at UC San Diego.

#### 5. Sustained guidance

To ensure the lasting success of BUMMP, an advisory board, consisting of the program’s founding members, has been established. The primary role of the advisory board is to provide oversight, ensuring that the leadership team consistently upholds the core values and mission of BUMMP. This becomes especially crucial with leadership turnover over time, assuring sustained guidance and alignment with the program’s objectives.

## Discussion

To combat the declining climate that is threatening DEI efforts within academia, it is imperative to intensify our endeavors in retaining students from HEGs. BUMMP was established with the goal of creating avenues of success tailored for first generation and historically underserved Biology undergraduate and master’s students at UC San Diego. The founders envisioned a future where each student has a sense of belonging and is guided by a clear path towards their future aspirations. This was achieved through an expansive reach, which provided access to quality mentorship, research, and professional development opportunities for over 1,350 mentees to date. The overarching purpose of this paper was to understand how mentorship, professional development, and research experiences positively influenced undergraduate and master’s science self-efficacy, identity, and sense of belonging. Based on prior research, we hypothesized that these experiences would lead to favorable outcomes in science self-efficacy, scientific identity, and a sense of belonging in the scientific community. We confirmed this hypothesis by demonstrating that a minimum of two quarters of mentorship, participation in professional development workshops, and research involvement significantly contributes to the integration of students into the STEM community, which is a favorable indicator for STEM retention and persistence. These findings suggest students’ experience matters. Considering that student persistence in STEM is influenced by factors such as confidence and sense of belonging, the expanded model of BUMMP will have a significant impact on a large number of students. To scale-up efforts in retaining students, particularly those from HEGs, the founders advocate that every institution adopts a mentorship program like BUMMP. Therefore, future efforts are aimed at collaborating with other institutions in establishing a BUMMP mentorship program, ultimately fostering an unprecedented level of retention for diverse students in STEM.

## Acknowledgments

This work was supported by CZI Diversity Leadership grant and NSF CAREER grant 2047391 to S.E.N, and Packard Foundation grant to R.D. We express our gratitude to all board members, mentors, program coordinators along with the Dean’s Office, Development Team and Equity, Diversity, and Inclusion Committee in the School of Biological Sciences at UC San Diego for their steadfast dedication to and invaluable support of BUMMP.

## Data Availability Statement

The data that support the findings of this study are available on request from the corresponding author. The data are not publicly available due to privacy or ethical restrictions.

---

**Figure S1:**
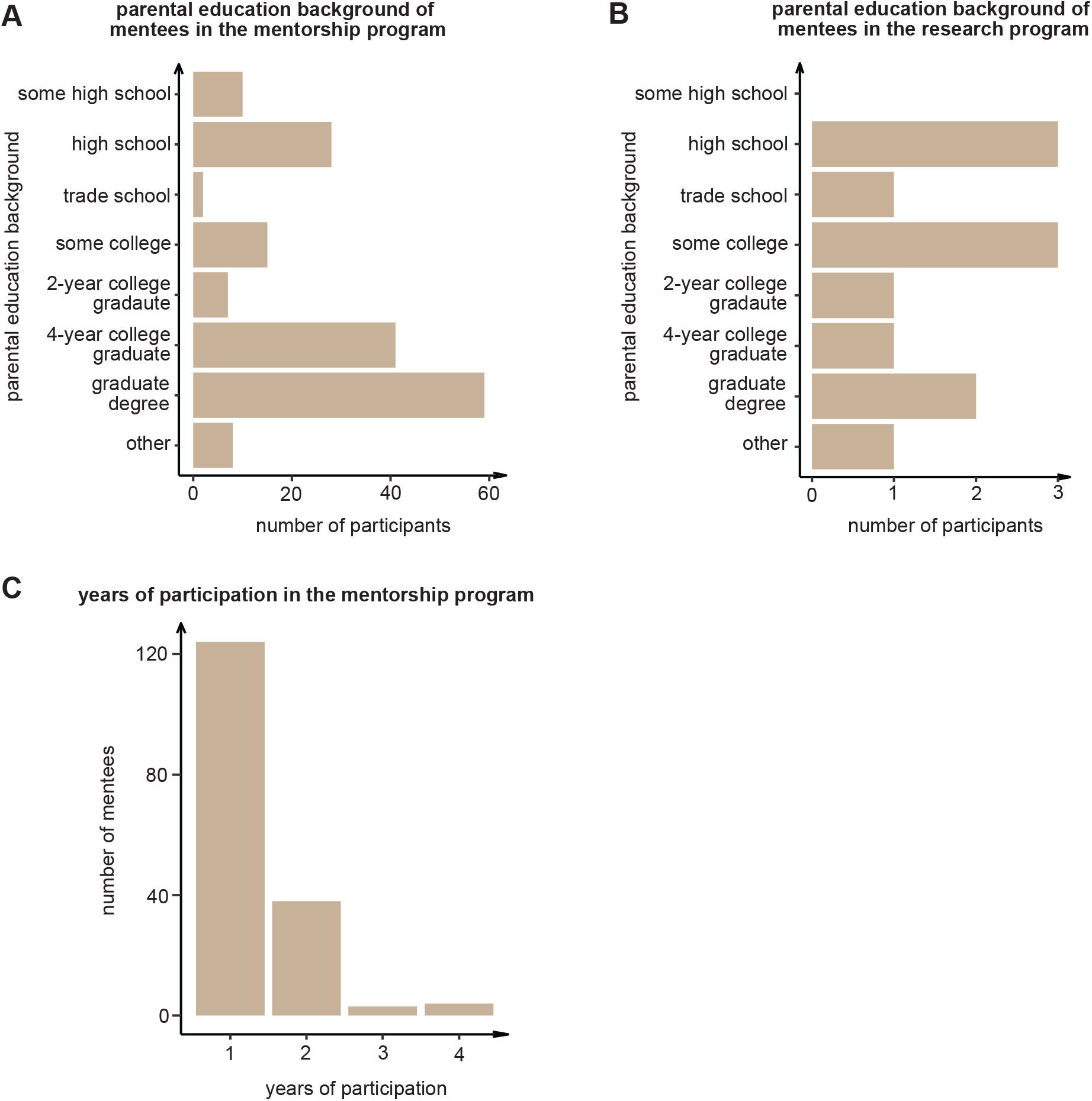
Backgrounds of mentees in the mentorship program and BRAP. (A) Highest level of education of the parent’s of mentees in the mentorship program. (B) Highest level of education of the parent’s of mentees in BRAP. (C) Total number of years a mentee has participated in the mentorship program.

**Figure S2:**
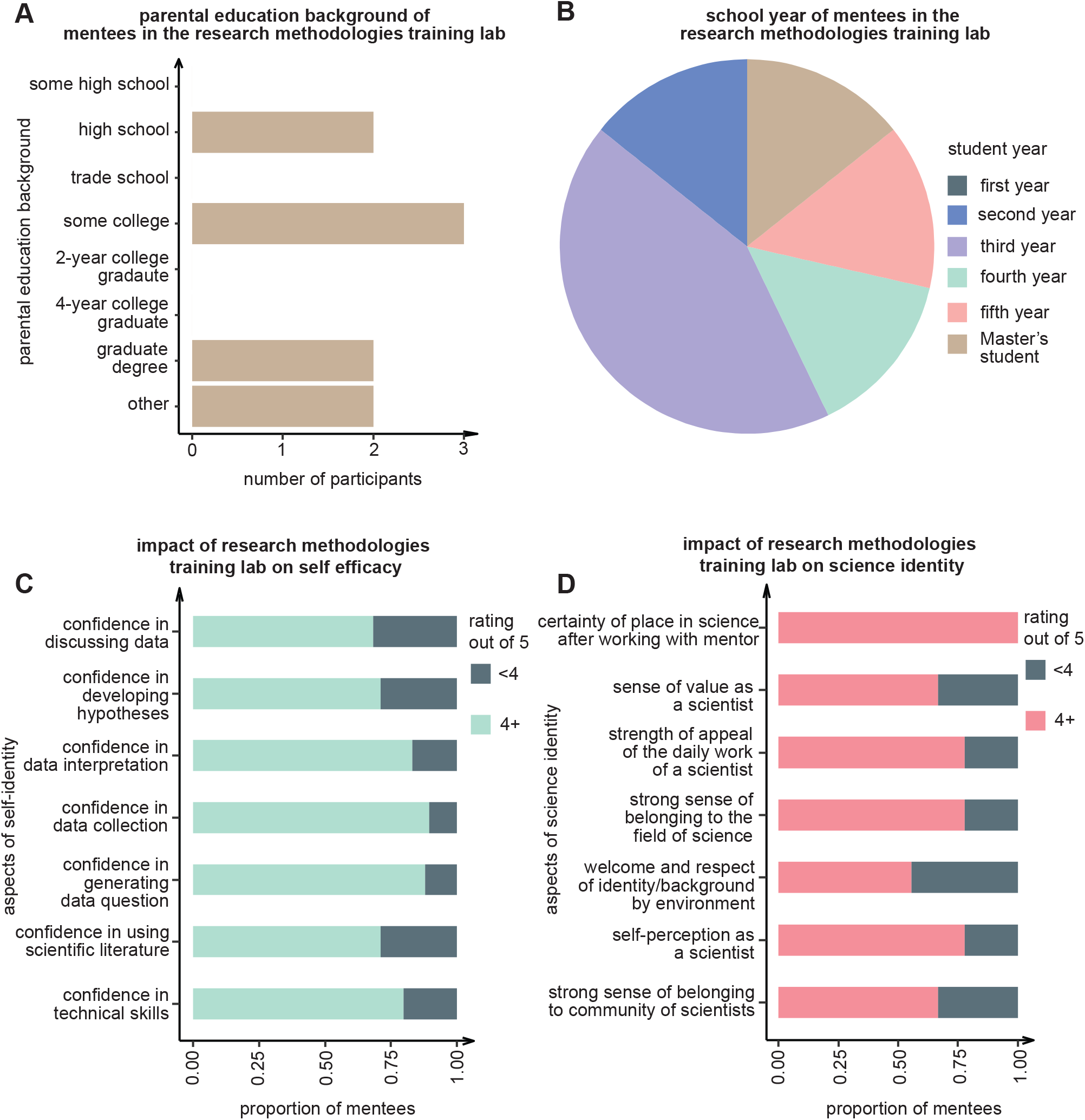
The research methodologies training lab, primarily composed of third years or higher, has a positive impact on science efficacy and identity. (A) Highest level of education of the parent’s of mentees in the research methodologies training lab (RMTL). (B) School year of mentee enrolled in the RMTL at the time of participation. (C) Impact of the RMTL on mentee’s self-efficacy. Mentees self-ranking on a scale of 1 to 5, 5 being the highest. (D) Impact of the RMTL on mentee’s science identity. Mentees self-ranking on a scale of 1 to 5, 5 being the highest.

## References

1. Berthoud, D. F., Forbes, D., & Garrett, F. (2021). *UC San Diego Black Academic Excellence Initiative.* https://diversity.ucsd.edu/initiatives/black-academic-excellence/BAEI-Recommendations-Summary-Report-2021---FINAL.pdf

2. Bryant, R. (2015). *College Preparation for African Americann Students: Gaps in the High School Educational Experience.* https://files.eric.ed.gov/fulltext/ED561728.pdf

3. Blau G., Surges D., Kory T., Ward-Cook, K., & Guiles H. (2003). Correlates of Fundamental Skills Versus Complex Skills for Medical Technologists. J. Allied Health, 32(1) https://www.jstor.org/stable/48721164

4. Estrada, M., Hernandez, P. R., & Schultz, P. W. (2018). A Longitudinal Study of How Quality Mentorship and Research Experience Integrate Underrepresented Minorities into STEM Careers. CBE Life Sciences Education, 17(1). 10.1187/CBE.17-04-0066

5. Gladstone, J. R., & Cimpian, A. (2021). Which role models are effective for which students? A systematic review and four recommendations for maximizing the effectiveness of role models in STEM. International Journal of STEM Education, 8(1). 10.1186/S40594-021-00315-X

6. Gopalan, M., Linden-Carmichael, A., & Lanza, S. (2022). College Students’ Sense of Belonging and Mental Health Amidst the COVID-19 Pandemic. Journal of Adolescent Health, 70(2), 228–233. 10.1016/j.jadohealth.2021.10.010

7. Haeger, H., Bueno, E. H., & Sedlacek, Q. (2024). Participation in Undergraduate Research Reduces Equity Gaps in STEM Graduation Rates. CBE Life Sciences Education, 23(1). 10.1187/CBE.22-03-0061/ASSET/IMAGES/LARGE/CBE-23-AR11-G003.JPEG

8. Haeger, H., & Fresquez, C. (2016). Mentoring for Inclusion: The Impact of Mentoring on Undergraduate Researchers in the Sciences. CBE Life Sciences Education, 15(3). 10.1187/CBE.16-01-0016

9. Hanauer, D. I., Graham, M. J., & Hatfull, G. F. (2016). A measure of college student persistence in the sciences (PITS). CBE Life Sciences Education, 15(4). 10.1187/CBE.15-09-0185/ASSET/IMAGES/LARGE/AR54FIG2.JPEG

10. Hanauer, D. I., & Hatfull, G. (2015). Measuring networking as an outcome variable in undergraduate research experiences. CBE Life Sciences Education, 14(4). 10.1187/CBE.15-03-0061/ASSET/IMAGES/LARGE/AR38FIG3.JPEG

11. Hathaway Biren, A., Nagda Sandra, R., Gregerman, R. S. (2002). The Relationship of Undergraduate Research Participation to Graduate and Professional Education Pursuit: An Empirical Study. 43(1).

12. Herres, J., Ortelli, O., Rodriguez, I., & Onyewuenyi, A. C. (2023). Factors associated with perceived stress and depressive symptoms among college students during the COVID-19 pandemic. Journal of American College Health: J of ACH. 10.1080/07448481.2023.2266039

13. Kim, E. (2009). Navigating College Life: The Role of Peer Networks in First-Year College Adaptation Experience of Minority Immigrant Students. Journal of the First-Year Experience & Students in Transition.

14. Lewis-McCoy, R. L. (2020). Inequality in the Promised Land. Inequality in the Promised Land. 10.1515/9780804792455/HTML

15. Lindstrom, L., Lind, J., Beno, C., Gee, K. A., & Hirano, K. (n.d.). Career and College Readiness for Underserved Youth: Educator and Youth Perspectives. Youth & Society, 2022(2), 221–239. 10.1177/0044118X20977004

16. Lopatto, D. (2007). Undergraduate research experiences support science career decisions and active learning. CBE Life Sciences Education, 6(4), 297–306. 10.1187/CBE.07-06-0039/ASSET/IMAGES/LARGE/CBE0040701010003.JPEG

17. Mariano, M. R., Sharp, S., Freeman, A., Freeman, T., Harmon, K., Wiggs, M., Sathy, V., Panter, A. T., Oseguera, L., Sun, S., Williams, M. E., Templeton, J., Folt, C. L., Barron, E. J., Hrabowski, F. A., Maton, K. I., Crimmins, M., Fisher, C. R., & Summers, M. F. (2019). Replicating Meyerhoff for inclusive excellence in STEM. Science, 364(6438), 335–337.

18. Marshall, A. G., Vue, Z., Palavicino-Maggio, C. B., Neikirk, K., Beasley, H. K., Garza-Lopez, E., Murray, S. A., Martinez, D., Crabtree, A., Conley, Z. C., Vang, L., Davis, J. S., Powell-Roach, K. L., Campbell, S., Brady, L. J., Dal, A. B., Shao, B., Alexander, S., Vang, N., … Hinton, A. (2022). The role of mentoring in promoting diversity equity and inclusion in STEM Education and Research. Pathogens and Disease, 80(1). 10.1093/FEMSPD/FTAC019

19. Noller, D. T., Bañuelas, A., & Diallo, S. (2024). Changing Course: Understanding the Legal Landscape of Race-Conscious Admission Practices and Implications to Diversity-Promoting Strategies in Higher Education. The Journal of Physician Assistant Education: The Official Journal of the Physician Assistant Education Association, 35(1), 101–104. 10.1097/JPA.0000000000000573

20. Pfund, C., Sancheznieto, F., Byars-Winston, A., Zárate, S., Black, S., Birren, B., Rogers, J., & Asai, D. J. (2022). Evaluation of a Culturally Responsive Mentorship Education Program for the Advisers of Howard Hughes Medical Institute Gilliam Program Graduate Students. CBE Life Sciences Education, 21(3). 10.1187/CBE.21-11-0321/ASSET/IMAGES/LARGE/CBE-21-AR50-G006.JPEG

21. Pierszalowski, S., Bouwma-gearhart, J., & Marlow, L. (2021). A Systematic Review of Barriers to Accessing Undergraduate Research for STEM Students: Problematizing Under-Researched Factors for Students of Color. Social Sciences 2021, Vol. 10, Page 328, 10(9), 328. 10.3390/SOCSCI10090328

22. Reeves, A. G., Bischoff, A. J., Yates, B., Brauer, D. D., & Baranger, A. M. (2023). A Pilot Graduate Student-Led Near-Peer Mentorship Program for Transfer Students Provides a Supportive Network at an R1 Institution. Journal of Chemical Education, 100(1), 134–142.

23. Riegle-Crumb, C., King, B., & Irizarry, Y. (2019). Does STEM Stand Out? Examining Racial/Ethnic Gaps in Persistence Across Postsecondary Fields. Educational Researcher, 48(3), 133–144.

24. Rinderknecht, F. A. B., Kouyate, A., Teklu, S., & Hahn, M. (2023). Antiracism in Action: Development and Outcomes of a Mentorship Program for Premedical Students Who Are Underrepresented or Historically Excluded in Medicine. Preventing Chronic Disease, 20. 10.5888/PCD20.220362

25. Robinson, G. F. W. B., Schwartz, L. S., Dimeglio, L. A., Ahluwalia, J. S., & Gabrilove, J. L. (2016). Understanding Career Success and Its Contributing Factors for Clinical and Translational Investigators. Academic Medicine : Journal of the Association of American Medical Colleges, 91(4), 570. 10.1097/ACM.0000000000000979

26. Romney, C. A., & Grosovsky, A. J. (2023). Mentoring to enhance diversity in STEM and STEM-intensive health professions. International Journal of Radiation Biology, 99(6), 983– 989. 10.1080/09553002.2021.1988182

27. Russell, S. H., Hancock, M. P., & McCullough, J. (2007). Benefits of undergraduate research experiences. Science, 316(5824), 548–549. 10.1126/SCIENCE.1140384/SUPPL_FILE/RUSSELL.SOM.PDF

28. Stebleton, M. J., & Soria, K. M. (n.d.). *Breaking down barriers: Academic obstacles of first-generation students at research universities*.

29. Tedesco, L. A. (2005). Post-Affirmative Action Supreme Court Decision: New Challenges for Academic Institutions. Journal of Dental Education, 69(11), 1212–1221. 10.1002/j.0022-0337.2005.69.11.tb04020.x

30. Tello, C., & Goode, C. A. (2023). Factors and barriers that influence the matriculation of underrepresented students in medicine. Frontiers in Psychology, 14. 10.3389/FPSYG.2023.1141045/FULL

31. Thiry, H., Laursen, S. L., & Hunter, A.-B. (2011). What Experiences Help Students Become Scientists? A Comparative Study of Research and other Sources of Personal and Professional Gains for STEM Undergraduates. The Journal of Higher Education, 82(4), 357–388. 10.1080/00221546.2011.11777209

32. *UCUES 2016 UC Systemwide Campus Report Very often Never Rarely Occasionally Somewhat often Often*. (n.d.).

